# Apixaban, an orally available anticoagulant, inhibits SARS-CoV-2 replication by targeting its major protease in a non-competitive way

**DOI:** 10.1101/2021.09.23.461605

**Authors:** Otávio Augusto Chaves, Carolina Q. Sacramento, Natalia Fintelman-Rodrigues, Jairo Ramos Temerozo, Filipe Pereira-Dutra, Daniella M. Mizurini, Robson Q. Monteiro, Leonardo Vazquez, Patricia T. Bozza, Hugo Caire Castro-Faria-Neto, Thiago Moreno L. Souza

**Affiliations:** Laboratory of Immunopharmacology, Oswaldo Cruz Institute (IOC), Oswaldo Cruz Foundation (Fiocruz), Rio de Janeiro, RJ, Brazil; National Institute for Science and Technology on Innovation on Neglected Diseases (INCT/IDN), Center for Technological Development in Health (CDTS), Oswaldo Cruz Foundation (Fiocruz), Rio de Janeiro, RJ, Brazil; National Institute for Science and Technology on Neuroimmunomodulation (INCT/NIM), Oswaldo Cruz Institute (IOC), Oswaldo Cruz Foundation (Fiocruz), Rio de Janeiro, RJ, Brazil; Laboratory on Thymus Research, Oswaldo Cruz Institute (IOC), Oswaldo Cruz Foundation (Fiocruz), Rio de Janeiro, RJ, Brazil; Institute of Medical Biochemistry Leopoldo de Meis, Federal University of Rio de Janeiro (UFRJ), Rio de Janeiro, RJ, Brazil

**Keywords:** SARS-CoV-2, apixaban, rivaroxaban, dabigatran, proteases, repurposing drugs

## Abstract

Anticoagulants are associated with clinical benefit against the 2019 coronavirus disease (COVID-19), preventing COVID-19 associated coagulopathy. Blood coagulation factor Xa (FXa) and SARS-CoV-2 major protease (M^pro^) share over 80% homology at the three-dimensional protein level. Thus, it is worth interrogating whether there is crosstalk between inhibitors and substrates between these enzymes. Here, we found that the clinically-approved FXa inhibitor apixaban targets SARS-CoV-2 M^pro^ with a 21-fold higher potency than boceprevir (GC376). Apixaban displayed a non-competitive mechanism of inhibition towards M^pro^, since it targets the enzyme/substrate complex and the allosteric site onto the viral protease. Enzymatic assays were further validated in infected Calu-3 cells, which reveal that apixaban decreases the production of infectious viral particles in a dose-dependent manner, with an inhibitory potency in the micromolar range. Our results are in line with the proposed early use of anticoagulants, including FXa inhibitors, to improve clinical outcome of COVID-19 patients. In this context, apixaban may display a dual mechanism of action by targeting FXa to prevent coagulopathy and, at some level, SARS-CoV-2 M^pro^.

## Introduction

The severe acute respiratory syndrome coronavirus 2 (SARS-CoV-2), the etiological agent of 2019 coronavirus disease 2019 (COVID-19), causes asymptomatic to life-threatening conditions, leaving 4.5 million deaths globally from January 2020 to August 2021^1,2^. Before reaching the severity of disease – characterized by acute respiratory failure, hyper-inflammation, and coagulopathy – early increase in blood levels of C reactive protein (CRP) and D-dimer already suggest poor clinical progression in COVID-19 patients^3,4^.

High D-dimer levels in COVID-19 patients are the final product of the hyperactivated clotting/fibrinolysis pathways and coagulation factor Xa (FXa) engagement is a key and rate-limiting event along with thrombin and fibrin generation^5,6^. These vascular changes in COVID-19 patients are enhanced by the pro-inflammatory cytokine storm that increases vascular permeability^7^. In fact, both cytokine storm and coagulopathy are triggered by the virus, as exemplified by monocytes from critically ill COVID-19 patients that expose tissue factor (CD142), a clotting trigger^8^. SARS-CoV-2 replication in type II pneumocytes leads to diffuse alveolar damage, and along with the formation of fibrin, documented in the necropsy of COVID-19 patients^5,9^, which may reduce alveolar hematosis.

Since the SARS-CoV-2 outbreak, the use of anticoagulants became part of the standard care in the management of COVID-19 patients^10-13^. Either broad and specific anticoagulants such as low-molecular weight heparin, direct thrombin/FXa inhibitors, and warfarin were recommended in the 2021 guidelines by both the American Society of Hematology (ASH) and National Institutes of Health (NIH) to critical and hospitalized patients^14-18^. Nevertheless, the posology of anticoagulants for COVID-19 patients may require adjustments; because current doses and regimens were imported from general practice to treat thrombosis and/or embolism^19^. Under these recently adapted conditions, antithrombotic treatment has been associated with decreased mortality in COVID-19 patients^16,18,20,21^; thus, confirming coagulopathy is a central event in the physiopathology of COVID-19.

The main protease (M^pro^) of SARS-CoV-2 is considered as one of the main targets for drug repurposing, due to its cleavage activity at eleven sites at the viral polyprotein^22^. Curiously, FXa and thrombin share a considerable similarity to SARS-CoV-2 M^pro^ binding site (according to the respective dimensionless scores of 0.71 and 0.74, in a maximum of 1.00). Compared to SARS-CoV-2 M^pro^, FXa and thrombin superpose their three-dimensional structures with a minor differences, with respect to Root Mean Square Deviation (RMSD) analysis of just 2.57 and 2.49 Å, respectively^22,23^. Although structural similarities between FXa and thrombin with M^pro^ have been suggested, functional studies to indicate whether M^pro^ could use FXa and thrombin inhibitors or substrates are scarce. Here, we demonstrated that the direct FXa inhibitor apixaban blocks SARS-CoV-2 M^pro^ enzymatic activity in a non-competitive way. As a consequence, apixaban impairs SARS-CoV-2 replication in a dose-dependent manner in human lung epithelial (Calu-3) cells.

## Results

### Apixaban non-competitively inhibits SARS-CoV-2 M^pro^

The FXa and thrombin show considerable similarity to M^pro^ binding site, as judged by the 3D superimposition of their structures with a good correlation (Fig. 1A) and in line with previous literature^22,23^. Thus, we interrogated if the M^pro^ (and as a control papain-like protease, PL^pro^) was susceptible to direct inhibitors of FXa (apixaban and rivaroxaban) and thrombin (dabigatran). Among the anticoagulants tested, apixaban displayed Morrison’s inhibitory constant (K_i_) values consistent with inhibition of a good inhibitory profile for M^pro^, but not for PL^pro^ (Fig. 1B and C). No compound inhibited PL^pro^ better than the positive control GRL0617 (Fig. 1B). With respect to the M^pro^, apixaban was 21-fold more potent than the positive controls boceprevir (GC376) and atazanavir, both on clinical trials against COVID-19 (Fig. 1C). Of note, the FXa inhibitor, ribaroxaban and the thrombin inhibitor, dabigatran were as potent as boceprevir (Fig. 1C).

**Fig. 1.**
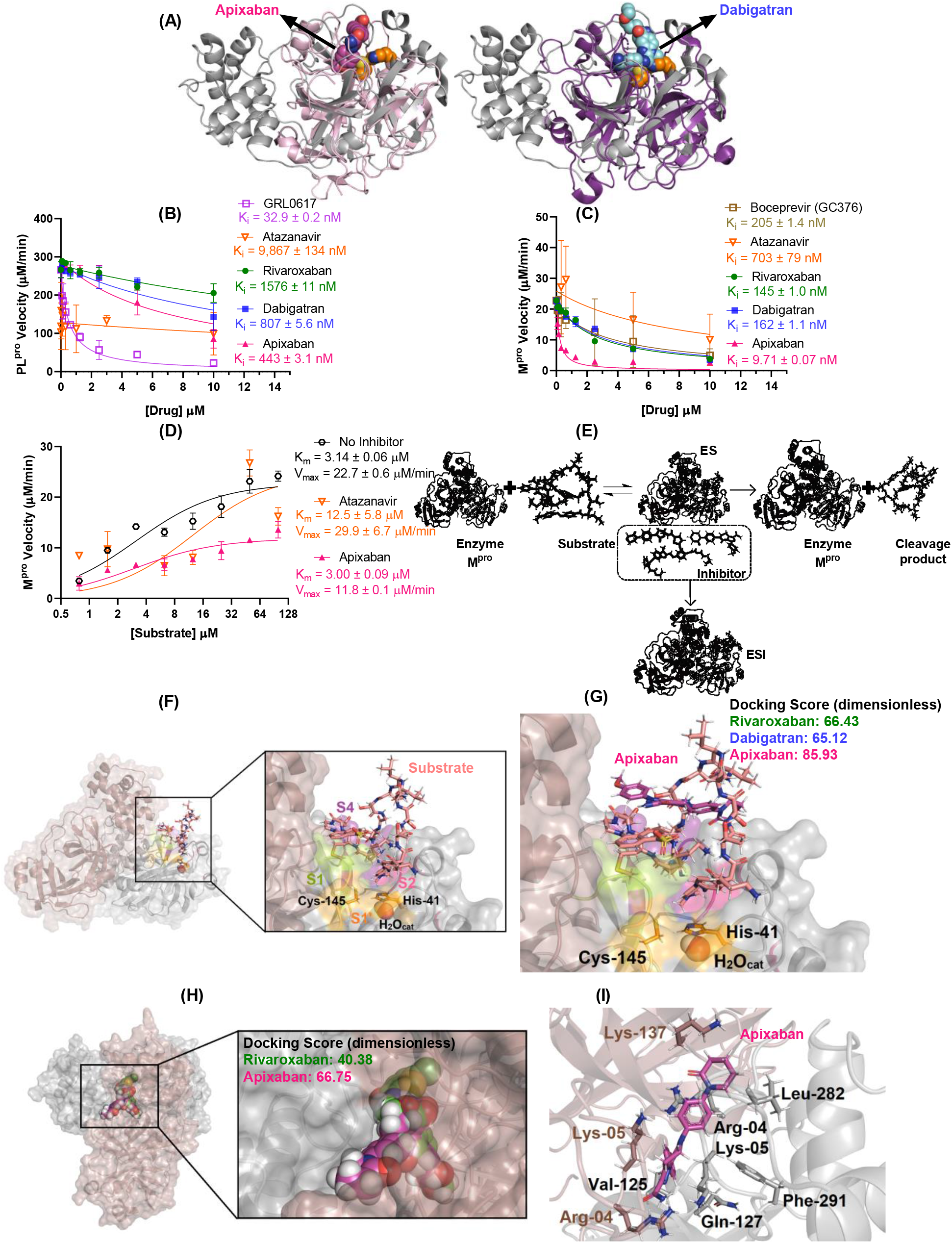
(A) Superposition of the monomeric unit of M^pro^ (in gray, PDB code 7K40) with FXa (on the left in violet, PDB code 2P16) and thrombin (on the right in purple, PDB code 1KTS). The crystallographic structure of apixaban into FXa structure, dabigatran into thrombin structure, and catalytic dyad of M^pro^ (His-41 and Cys-145 residues) are as spheres in pink, cyan, and orange, respectively. For better interpretation the catalytic water (H_2_O_cat_) of M^pro^ is not shown. The enzymatic inhibition profile for apixaban, rivaroxaban, and dabigatran (0, 0.08, 0.16, 0.31, 0.63, 1.25, 2.5, 5.0, and 10 mM) into (B) PL^pro^ (8.19 nM) and (C) M^pro^ (88.8 nM) velocity. The positive controls GRL0617 (PL^pro^) and GC376 (M^pro^) were used under the same condition of anticoagulants. (D) Michaelis-Menten enzymatic mechanism for M^pro^ without and in the presence of a fixed apixaban or atazanavir concentration (2.5 mM) for different substrate concentrations (0, 0.76, 1.56, 3.12, 6.25, 12.5, 25.0, 50.0, and 100 mM). (E) Enzymatic scheme for the experimental mechanism of M^pro^ inhibition by anticoagulants. Best docking pose (ChemPLP function) for the interaction between M^pro^ (F) substrate, and (G) substrate-apixaban into the active site of protease. Best docking pose (ChemPLP function) for the interaction between the dimer interface of M^pro^ (H) apixaban and rivaroxaban, while (I) shows the selected amino acid residues which interact with apixaban. Substrate, rivaroxaban, dabigatran, and apixaban are in stick representation in beige, green, cyan, and pink, respectively, while the catalytic water (H_2_O_cat_) is in sphere. Elements’ color: hydrogen, nitrogen, oxygen, sulfur, and chloro are in white, dark blue, red, yellow, and dark green, respectively.

Since apixaban inhibits M^pro^ with a K_i_ lower than the concentration of the viral protease used in the assay, a non-canonical mechanism of inhibition over this enzyme might be expected. When apixaban’s inhibition over M^pro^ was assayed under different concentrations of substrate, a non-competitive mechanism was observed (Fig. 1D). The Michaelis-Menten constant (K_m_) value was not altered by apixaban, suggesting M^pro^ is not halted to interact with its substrate by apixaban (Fig. 1D). In addition, there was a significant decrease in the M^pro^ maximum velocity (V_max_) by apixaban (Fig. 1D), indicating that the enzyme (E) M^pro^ bound to its substrate (S), in the enzyme-substrate complex (ES) and was unable to cleave it and form the products (P) (Fig. 1E).

Based on molecular docking, we explored different alternative to explain these results. When interacting with M^pro^, its peptidic substrate occupies the four enzymatic subsites (S1, S1’, S2, and S4) onto the active binding pocket with a docking score (dimensionless) of 65.54 (Fig. 1F). In the M^pro^/peptide (ES) complex, the substrate forms an external loop, which is targeted by the anticoagulants (Fig. 1G); in particular, apixaban has the most favorable interaction compared with dabigatran and rivaroxaban due to the significant hydrogen bonding interaction with substrate (3.00 Å, Fig. 1G and Supplementary Figs. 2A-C). Moreover, three additional binding pockets in the dimer interface of M^pro^ were identified and associated with allosteric regulation^24,25^. Molecular docking calculations suggested that apixaban binds onto the allosteric site of M^pro^ with high number of hydrogen bonding and van der Waals interactions (Figs. 1H and 1I). Rivaroxaban did not reach the allosteric site; just the M^pro^/M^pro^ dimeric interface and dabigatran did not present a feasible binding capacity into these additional sites (Fig. 1H and Supplementary Fig. 2D).

Next, we evaluated if whether SARS-CoV-2 proteases could cleave the synthetic chromogenic substrates designed for thrombin (S-2238) and coagulation FXa (S-2765). Despite M^pro^ and FXa similarities described above, the substrates were not cleaved by M^pro^, nor by PL^pro^, also included as control (Supplementary Fig. 1A). In fact, host and viral enzymes belong to different families of endopeptidases and M^pro^ lacks the superimposed of random coils that are external of the active site of FXa, which might impact the mimetic chromogenic substrate accommodation into M^pro^ active site (Fig. 1A).

### Apixaban inhibits SARS-CoV-2 replication in a dose-dependent manner in pneumocyte cell lineage

To further demonstrate SARS-CoV-2 susceptibility to anticoagulants, Calu-3 that recapitulates the most affected cells in the respiratory tract, the type II pneumocytes^26^, were infected and treated with these compounds. Anticoagulants inhibited the production of infectious SARS-CoV-2 progeny in a dose-dependent manner (Figs. 2A and 2B). Consistently with the enzymatic data, apixaban was approximately 3-fold more potent than any other anti-clotting drugs tested (Table 1). Nevertheless, apixaban was about 5- and 60-fold less potent in vitro in comparison to the positive control atazanavir and remdesivir, respectively (Table 1), indicating that apixaban shows an interestingly scaffold for the design of novel compounds to increase its antiviral action. Despite a slightly higher cytotoxicity, compared to other tested compounds, apixaban’s selectivity index (SI) for SARS-CoV-2 replication was two-fold better than other anticoagulants (Table 1). Altogether our results confirm that apixaban’s chemical structure is endowed with antiviral activity against SARS-CoV-2.

**Table 1.** The 50% effective concentration (EC_50_ for MOI 0.1), 50% cytotoxic concentration (CC_50_), and selectivity index (SI) of the anticoagulants, atazanavir, and remdesivir in Calu-3 cells.

| Anticoagulants | $EC_{50}$ (μM) | $CC_{50}$ (μM) | SI |
| --- | --- | --- | --- |
| Atazanavir | $0.353 \pm 0.018$ | $312 \pm 8$ | 884 |
| Remdesivir | $0.0305 \pm 0.0031$ | $512 \pm 30$ | $1.68 \times 10^4$ |
| Rivaroxaban | $5.90 \pm 0.30$ | $553 \pm 28$ | 93.7 |
| Dabigatran | $6.94 \pm 0.35$ | $756 \pm 38$ | 109 |
| Apixaban | $1.84 \pm 0.09$ | $491 \pm 25$ | 267 |

**Fig. 2.**
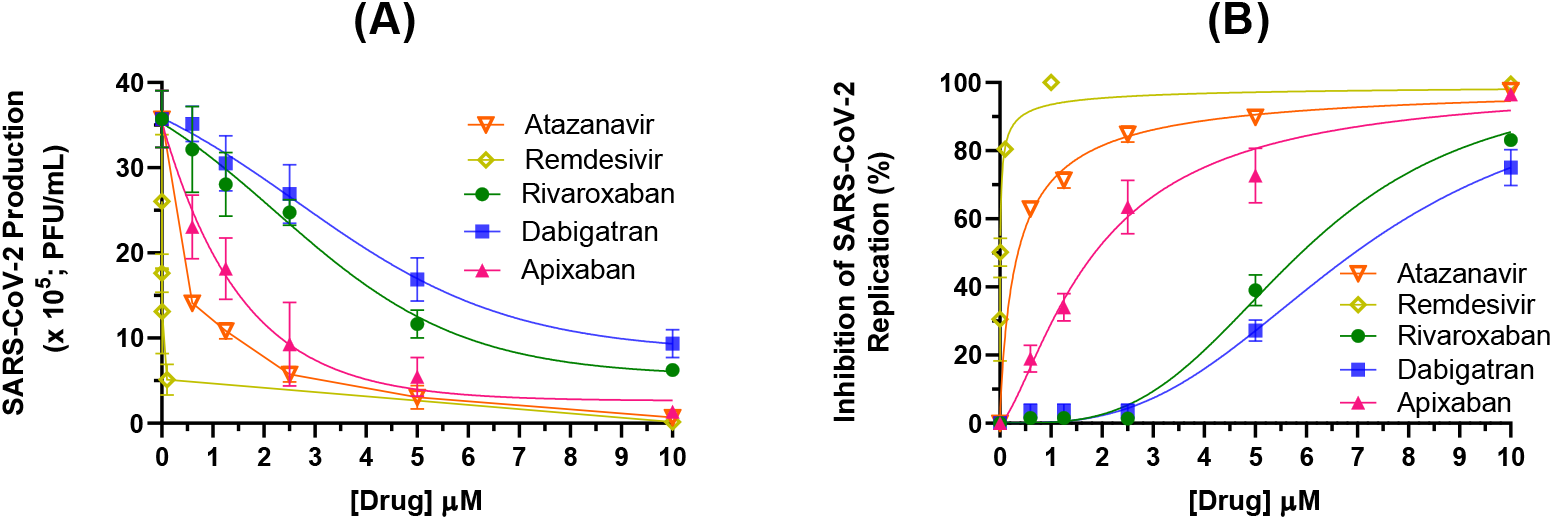
Antiviral activity of anticoagulants, atazanavir, and remdesivir in Calu-3 cells (densities of 2.0 × 10^5^ cells/well) infected with SARS-CoV-2 (MOI 0.1) in 96-well plates. The data is presented as (A) virus production (PFU/mL) and (B) percentage of viral replication inhibition. The data represent means ± SEM of three independent experiments.

## Discussion

Severe COVID-19 could be characterized as a SARS-CoV-2-triggered host-related cytokine storm and coagulopathy^27^. In this sense, clinical trials have been investigating whether COVID-19 patients could benefit from oral anticoagulant agents, such FXa and thrombin inhibitors^14-21^. Indeed, an increasing body of evidence reinforces that anti-clotting drugs increase the survival probability of critically ill COVID-19 patients under non- and invasive ventilatory assistance^15,16,18,21,28^. Moreover, other clinical studies also suggest that the early use of anti-thrombotic agents could prevent the consumption of platelets, clotting factors, and ultimately hemorrhagy^16,19,28^. The high levels of D-dimer, a final product from clotting and fibrinolysis, directly implicate these cascades in the physiopathology of COVID-19^28^. Thus, the benefit from anticoagulants is well rationalized as host-acting agents. Nevertheless, because of the structural similarities between SARS-CoV-2 M^pro^ and FXa^23^, we studied here if: i) M^pro^ could directly cleave FXa substrate, for comparisons we also included PL^pro^ and other substrates in the analysis; ii) FXa inhibitors could also directly target viral M^pro^ and replication, for comparison PL^pro^ and other inhibitors were included. We found that apixaban is a potent M^pro^ inhibitor with a unique mechanism of action and, consequently, reduced SARS-CoV-2 replication.

Remarkably, the oral anti-factor Xa drug apixaban was a potent M^pro^ inhibitor, comparable to other repurposed drugs on clinical trials against COVID-19, such as the HIV protease inhibitor atazanavir. Differently from atazanavir, apixaban was a non-competitive inhibitor of M^pro^, targeting the allosteric site of the viral enzyme and its secondary complex bound to the substrate. These characteristics suggest that apixaban chemical structure could be even worth for further hit-to-lead development against COVID-19. The cell-free-based assays were further validated in SARS-CoV-2 replicative experiments, confirming that apixaban directly inhibits SARS-CoV-2 replication.

Under clinically approved posology of 10 mg, apixaban reaches a maximum plasmatic concentration (C_max_) of 0.55 μM^29^. Considering that 87% of the apixaban is bound to albumin^30^, its free fraction at C_max_ is equivalent to 72 nM, almost 10-times higher than apixaban’s K_i_ towards M^pro^. On the other side, cell-based assays, display apixaban’s potency as 3-times higher than human C_max_. Despite apixaban is more potent than atazanavir to inhibit M^pro^, the IC_50_ value for apixaban was not better than atazanavir, probably due to the influence of permeability into the cells: logP for apixaban and atazanavir of 2.71 and 4.50, respectively. These comparisons, of cell-free and -based experiments on the pharmacokinetic parameters lead to paradoxical interpretations on whether apixaban could be acting as a direct acting antivirals in COVID-19 patients. We interpret that cell-based assay conditions are subjected of greater interference than enzymatic experiments. For example, Calu-3 infection was conducted at the considerable multiplicity of infection of 0.1 (1 virus plaque forming unit/10 cells) and cells are maintained with 10% fetal bovine serum, which may bind apixaban. Enzyme kinetic assays were suggestive of a direct acting antiviral activity – which is in line with early clinical use of apixaban during COVID-19^31,32^.

It is naturally difficult to estimate a clinical benefit of any antiviral activity of apixaban during clinical trials because its anti-clotting activity is directly associated with COVID-19 physiopathology. Nevertheless, apixaban chemical structure could even be optimized to the development of novel non-competitive M^pro^ inhibitors that preserve anticoagulant activity.

Finally, under our experimental conditions, M^pro^ was unable to cleave S-2765, a mimetic chromogenic substrate that recapitulates pro-thrombin cleavage site, possibly due to structural and functional differences between M^pro^ and FXa at the random coil region.

## Methods

### General

All reagents were purchased from Sigma-Aldrich/Merck (St. Louis, MO, USA), HyClone Laboratories Inc. (Logan, Utah) or Chromogenix (Diapharma Group, Inc., KY). The SARS-CoV-2 virus was isolated from a nasopharyngeal swab of a confirmed case from Rio de Janeiro, Brazil, and its complete genome was sequenced and publicly deposited (GenBank #MT710714; Institutional Review Broad approval, 30650420.4.1001.0008).

### Enzymatic assays

The anticoagulant capacity in inhibit enzymatic activity of M^pro^ and papain-like protease (PL^pro^) from SARS-CoV-2 were determined by the commercial kit provided by BPS Biosciensces® company (catalog number: #79955-1 and #79995-1, respectively) following the procedure and recommendations from literature and company^33,34^. The enzymatic inhibition was evaluated by varying concentrations of the anti-clotting drugs. Next, the mechanism of inhibition was determined via Michaelis-Menten by varying substrate concentration.

The M^pro^ and PL^pro^ were assayed for amidolytic activity towards the hydrolysis of the synthetic chromogenic substrates for thrombin (S-2238) and coagulation factor Xa (S-2765). Hydrolysis of S-2238 and S-2765 (0.2 mM final concentration) by M^pro^ (10 nM) and PL^pro^ (10 nM) was determined in 50 mM Tris/HCl, 150 mM NaCl, 10 mM CaCl_2_, and 0.1% polyethylene glycol (PEG) 6000, pH 7.5. Substrate hydrolysis was detected using a SpectraMax® ABS Plus equipped with a microplate mixer and heating system. Reactions were recorded continuously at 405 nm for 2 hours at 37 °C.

### In vitro assays

The Calu-3 cells were infected with multiplicity of infection (MOI) of 0.1 at densities of 2.0 × 10^5^ cells/well for 1 h at 310K in 5 % of CO_2_. The cells were washed, and different concentrations of each anticoagulant, atazanavir, or remdesivir were added in Dulbecco’s modified Eagle medium (DMEM) with 10% fetal bovine serum (FBS). The concentration for anticoagulants and atazanavir was 0.00, 0.63, 1.25, 2.50, 5.00, and 10.0 μM, while remdesivir was 0.00, 0.0001, 0.001, 0.01, 0.10, 0.50, 1.0, 5.0, and 10.0, μM. After 48 h, the supernatants were harvested, and virus titers were quantified by plaque-based assays according to previous publications^36-38^,

The cytotoxic assays were conducted in a monolayers of Vero cells (in about 2.0 × 10^4^ cell/well) treated for 3 days with different concentrations of apixaban, rivaroxaban, dabigatran, atazanavir, or remdesivir (50, 150, 300, 600, and 800 µM) following procedure described by Sacramento, C.Q. et al^37^. The plates were read in terms of absorption in a spectrophotometer at 595 nm and the 50% cytotoxic concentration (CC_50_) was calculated by a non-linear regression analysis from a dose– response curve.

### Statistics

All in vitro data were analyzed from Prism GraphPad software 8.0 (Windows GraphPad Software, San Diego, California USA). At least triplicate experiments were performed for each data point, and the value was presented as mean ± standard deviation (SD).

### Molecular docking procedure

The crystallographic structure for M^pro^ was obtained from Protein Data Bank, with access code 7K40. The chemical structure for the anticoagulants and M^pro^ substrate used in the experimental assays was built and minimized in terms of energy by Density Functional Theory (DFT) via Spartan’18 software (Wavefunction, Inc., Irvine, CA, USA). The molecular docking calculations were performed with GOLD 2020.2 software (Cambridge Crystallographic Data Center Software Ltd., CCDC) at pH 7.4. Redocking studies were carried out with the crystallographic ligand boceprevir (GC376), obtaining the lowest RMSD value by ChemPLP function. The main binding pockets for drugs into M^pro^ were calculated by the free access software from Proteins Plus (Zentrum für Bioinformatik, Universität Hamburg, Germany). It was defined 8 Å radius around each main binding pockets and the figures of the best results were generated with PyMOL Delano Scientific LLC software (DeLano Scientific LLC: San Carlos, CA, USA).

## Acknowledgements

The authors thank the Brazilian agencies for the financial support: Conselho Nacional de Desenvolvimento Científico e Tecnológico (CNPq), and Fundação Carlos Chagas Filho de Amparo à Pesquisa do Estado do Rio de Janeiro (FAPERJ). This study was financed in part by Coordenação de Aperfeiçoamento de Pessoal de Nível Superior (CAPES, Brazil) with finance code 001.

## Authors contribution

T.M.L.S., H.C.C.-F.-N., and P.T.B. idealized the work. O.A.C., C.Q.S., N.F.-R., and J.R.T. conducted the SARS-CoV-2 inhibition in Calu-3 cells. O.A.C., F.P.-D., and L.V. conducted the experimental proteases assays. O.A.C. conducted the cytotoxic assays and molecular docking calculations. D.M.M. and R.Q.M. conducted the pro-clotting coagulation assays. O.A.C., R.Q.M., P.T.B., H.C.C-F-N., and T.M.L.S. manuscript preparation.

## Competing interests

The authors declare no competing interests.

## Supplementary Information

**Supplementary Fig. 1.**
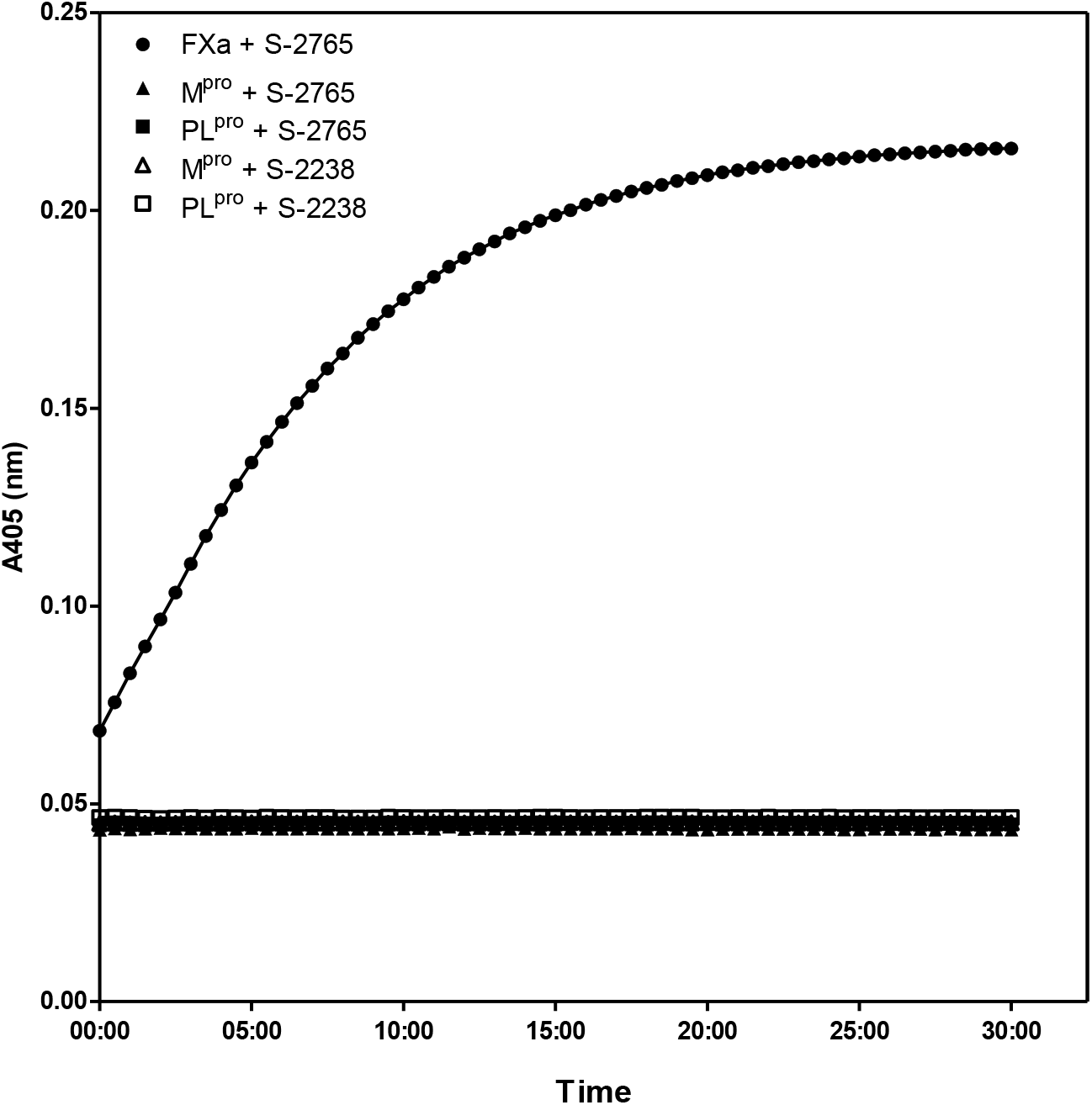
Cleavage of chromogenic substrates S-2765 and S-2238 by M^pro^ and PL^pro^. M^pro^ (10 nM) and PL^pro^ (10 nM) were incubated with S-2765 or S-2238 (0.2 mM final concentration) and the kinetic of the reactions was monitored by measuring the absorbance at 405 nm for thirty minutes at room temperature. (●) indicate control performed in the presence of FXa (1.25 nM). Results are expressed as mean values ± SEM of duplicates.

**Supplementary Fig. 2.**
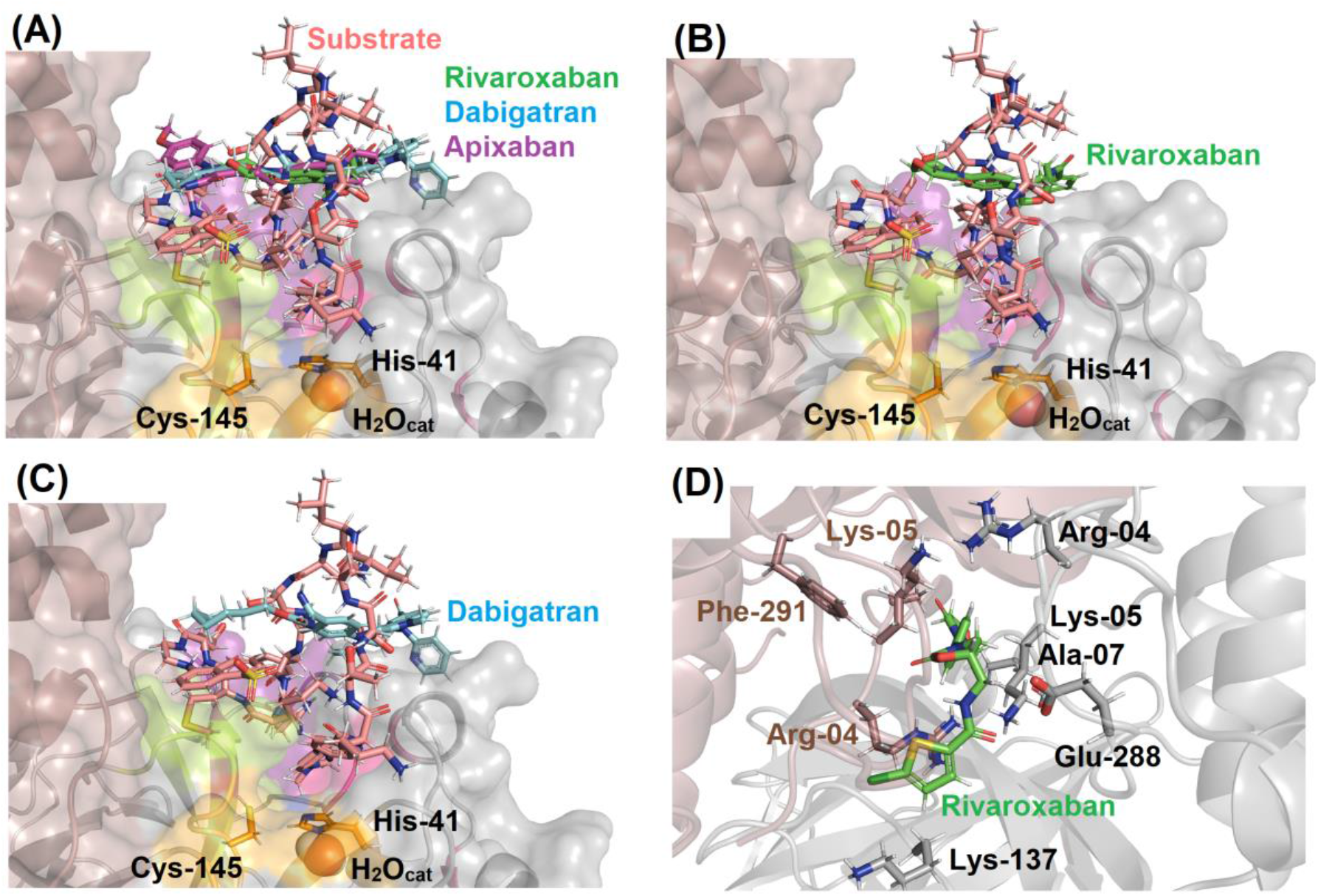
Best docking pose (ChemPLP function) for the interaction between M^pro^ (A) substrate-anticoagulants, (B) substrate-rivaroxaban, and (C) substrate-dabigatran into the active site of protease. (D) Best docking pose (ChemPLP function) for the interaction between rivaroxaban into the dimer interface M^pro^. Substrate, rivaroxaban, dabigatran, and apixaban are in stick representation in beige, green, cyan, and pink, respectively, while the catalytic water (H_2_O_cat_) is in sphere. Elements’ color: hydrogen, nitrogen, oxygen, sulfur, and chloro are in white, dark blue, red, yellow, and dark green, respectively.

## Notes

### Competing Interest Statement

The authors have declared no competing interest.

